# Projection of gut microbiome pre and post-bariatric surgery to predict surgery outcome

**DOI:** 10.1101/2020.08.27.271312

**Authors:** Meirav Ben Izhak, Adi Eshel, Ruti Cohen, Liora Madar Shapiro, Hamutal Meiri, Chaim Wachtel, Conrad Leung, Edward Messick, Narisra Jongkam, Eli Mavor, Shimon Sapozhnikov, Nitsan Maharshak, Subhi Abu-Abeid, Avishai Alis, Ilanit Mahler, Aviel Meoded, Shai Meron Eldar, Omry Koren, Yoram Louzoun

## Abstract

**Background:** Bariatric surgery is often the preferred method to resolve obesity and diabetes, with ~800,000 cases worldwide yearly and high outcome variability. The ability to predict the long-term Body Mass Index (BMI) change following surgery has important implications on individuals and the health care system in general. Given the tight connection between eating habits, sugar consumption, BMI, and the gut microbiome, we tested whether the microbiome before any treatment is associated with different treatment outcomes, as well as other intakes (high-density lipoproteins (HDL), Triglycerides, etc.).

**Results:** A projection of the gut microbiome composition of obese (sampled before and after bariatric surgery) and slim patients into principal components was performed and the relation between this projection and surgery outcome was studied. The projection reveals 3 different microbiome profiles belonging to slim, obese, and obese who underwent bariatric surgery, with post-surgery more different from the slim than the obese. The same projection allowed for a prediction of BMI loss following bariatric surgery, using only the pre-surgery microbiome.

**Conclusions:** The gut microbiome can be decomposed into main components depicting the patient’s development and predicting in advance the outcome. Those may be translated into better clinical management of obese individuals planning to undergo metabolic surgery.

**Importance:** BMI and diabetes can affect the gut microbiome composition.

Bariatric surgery has large variabilities in outcome.

The microbiome was previously shown to be a good predictor for multiple diseases.

We analyzed here the gut microbiome before and after bariatric surgery and show that:

- The microbiome before surgery can be used to predict surgery outcome.
- Post-surgery microbiome drifts further away from the slim microbiome than pre-surgery obese patients.

These results can lead to a microbiome-based pre-surgery decision whether to perform surgery.

## Background

The human body is colonized by a wide variety of micro-organisms, commonly referred to as the human microbiota. The gut microbiota is a complex ecosystem, which provides major functions to the host, such as regulation of metabolism, immune system modulation, and protection against pathogens (1, 2). The microbiome is strongly associated with weight and sugar consumption, and as such it serves as a proxy for nutrition and life habits and may also influence them. Such life habits may influence the total body mass and BMI in regular conditions, as well as after bariatric surgery.

Obesity and diabetes are world pandemics (3). Approximately 8-10% of the population develop complications of morbid obesity, (BMI>35), frequently coupled to some form of diabetes. According to the WHO, of the 57 million deaths in 2008 worldwide, 1.3 million were due to metabolic disorders, particularly those associated with obesity (3). Recently, the gut microbiome of obese individuals has been shown to differ from the microbiome of slim subjects (4). Nagpal et al. (5) suggested that some bacteria increase gut permeability and insulin resistance leading to obesity and diabetes. Experimental fecal transplants to mice demonstrated that transplantation of microbiome from obese individuals into slim mice turned them obese (6), showing the importance of the gut microbiome in regulating body weight. Opposite studies of turning obese mice into slim mice have not been successful but one study demonstrated that certain bacteria can prevent weight gain in mice (7).

The introduction of bariatric surgery as a method for losing weight is rapidly adopted as the most efficient method for weight loss and for reducing blood sugar levels (8), however, it has drawbacks, including a range of possible complications from nutrition deficiencies to occurrence of life-threatening conditions and a big diversity in the weight loss rate and its maintenance (9). A few studies have shown microbiome changes after bariatric surgery. A recent systematic review (10) summarized the finding of 9 human studies and 12 animal studies described an increase in the relative abundance of 4 major phyla: Proteobacteria, Fusobacteria, Bacteroidetes and Verrucomicrobia as opposed to a decrease in the phylum Firmicutes. The dominant genera which changed were *Faecalibacterium*, *Lactobacillus* and *Coprococcus*. An interesting finding was the increase in microbial diversity post-surgery (11). One of the mechanisms proposed for the effectiveness of bariatric surgeries is the changes in microbiome which influence the bile acids composition leading to metabolic improvement (12, 13). This can also occur the other way around, changes in bile acids, pH, and hormone levels lead to a change in the microbiome which affects energy homeostasis (14). However, the real potential of the microbiome as a tool not only for monitoring the procedure’s outcomes but rather predicting them in advance has not been explored. An attempt to study and test this potential can result in an essential tool which will assist in the decision whether to consult patient to perform such surgery.

## Methods

### Patients and regulation

Patients were enrolled in the obesity control/bariatric surgery clinics of four medical centers in Israel – Kaplan Medical Center (KMC), Rabin Medical Center (RMC), Tel Aviv Medical Center (TMC) and Poria Medical Center (PMC) during December 2015-November 2018. The ethics committees of each of the respective medical centers approved the study and its amendments and each patient and control signed written informed consent. Inclusion criteria were: ages 18-70, no antibiotic treatment in the two months before enrolment, no previous bariatric or major gut/stomach operation. One-third of the patients had diabetes (type 1 and type 2) and were treated with either insulin (Type 1) or other drugs (metformin or others).

Naturally slim controls were defined as BMI 19-25, and are healthy controls recruited from the same population. The controls had no diabetes (Hemoglobin A1C<5.0), BMI of 19-25, and had no major medical and endocrine complications. Obese individuals had BMI values of 35 and above, with and without active diabetes type 1 or 2 that is treated with medications.

Naturally slim control individuals gave one fecal, blood and urine sample. Obese/diabetic people gave five samples of each at the following time points: time of enrolment (group A), three weeks after low carbohydrate diet and immediately before the operation (Group B), 2 weeks (group C), 3 and 6 months after the operation (groups D&E, respectively). In all visits, the patients’ weight, blood and urine test results, medications and general health issues were noted. The obese group also provided weight values and blood test results at 1 and 1.5 years after the operation. At the time of analysis, not all patients completed their course of testing and evaluation.

Blood tests included the standard Complete Blood Count (CBC) test list and the blood biochemistry test following 12 hr fasting (of relevance here triglycerides, LDL, HDK, blood glucose). Also, patients provide samples for the standard hemoglobin A1 C (HbA1C) test.

### DNA extraction

Fecal samples were stored in Flora prep tubes (Admera Health, New Jersey, USA) with a proprietary bacterial DNA preservation media, allowing for storage at room temperature. The samples were brought into the NGS lab within 1-2 days after collection and were subjected to DNA extraction, purification and cleaning. DNA was extracted from the stool sample using the PowerSoil DNA extraction kit (MoBio, Carlsbad, CA) according to the manufacturer’s instructions. Purified DNA was used for PCR amplification of the variable V3 and V4 regions (Using MetaVx.2™ system developed with GENEWIZ (Plainfield, NJ)of the 16S rRNA gene as was previously described in US patent 9,745611, and by Caporaso et al (15). Amplicons were purified using AMPure magnetic beads (Beckman Coulter, Brea, CA) and subsequently quantified using Qubit dsDNA quantification kit (Invitrogen, Carlsbad, CA). Equimolar amounts of DNA from individual samples were pooled and sequenced using the Illumina MiSeq platform and V2 500 cycle kit.

### DNA Amplification

Briefly, the amplification method includes using multiple sets of overlapping primers and generation of multiple frame-shifted amplicons, which increase taxonomically diverse sequence amplification. The primers used in the present work are listed in Table 1 and are designed to amplify the variable regions V3 and V4 of the bacterial 16S rRNA.

**Table 1:**
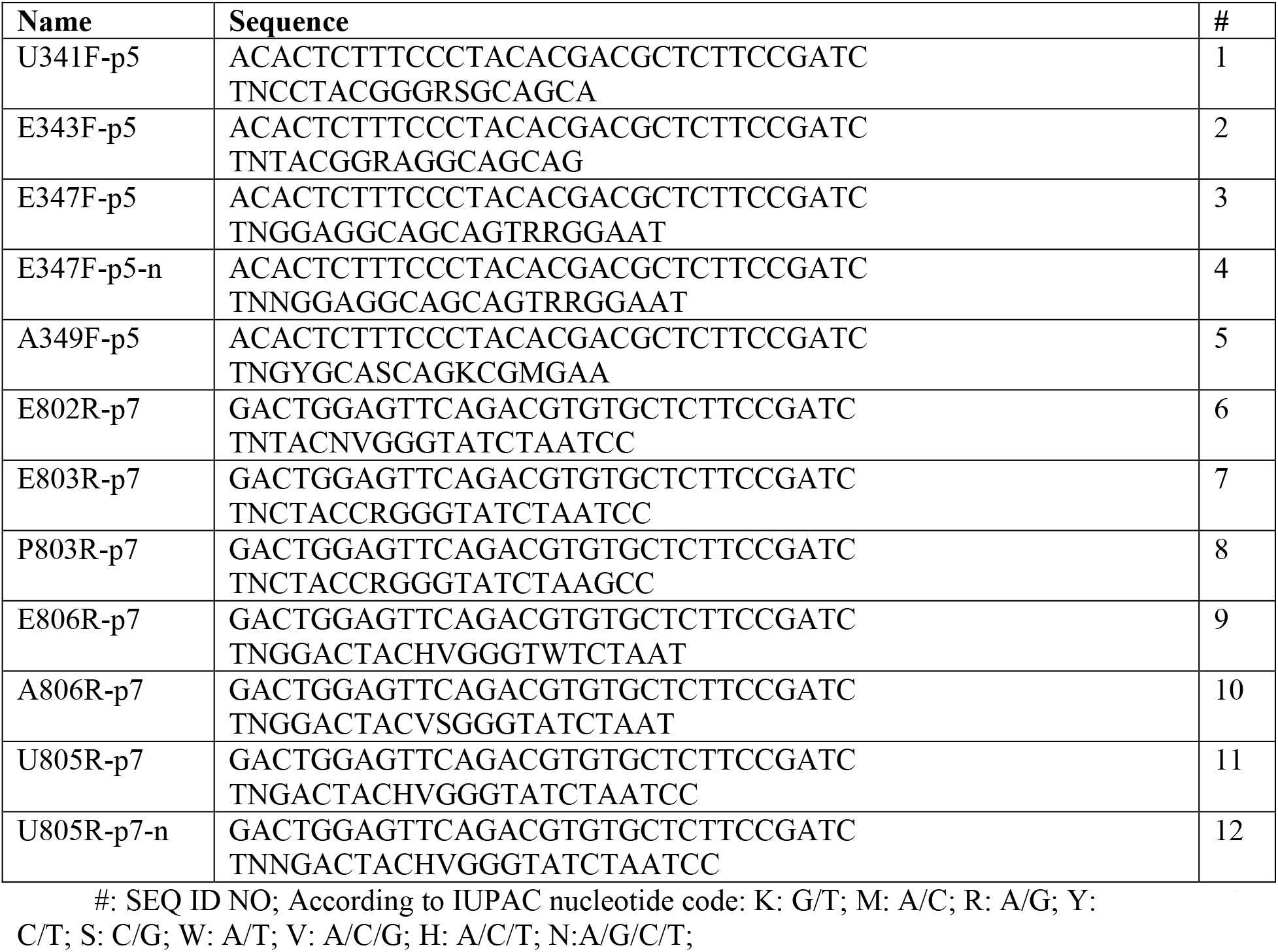
Primers sequences.

As can be seen from Table 1 of US patent 9,745,611, these primers are capable of amplifying sequences from bacterial gut microbiome to identify rare species which may be difficult to amplify by similar kits based on 16S rRNA V4 region sequence amplification. The use of this procedure enabled the identification of bacterial taxonomic entities below 0.2% of the population and the detection of rare species.

### Microbiome analysis

Microbial communities were analyzed using Quantitative Insights Into Microbial Ecology software QIIME2 (16). Single-end sequences were demultiplexed by per-sample barcodes and error-corrected by Divisive Amplicon Denoising Algorithm (DADA2) (17), primers were trimmed off and single-end reads were truncated to ≥ 160 base pairs. Feature sequences were aligned against Greengenes database v13_8 (18) with a similarity of 99% or greater, for taxonomic annotation. Finally, the following contaminants have been removed from the feature table: Thermi, S24-7, and Chloroplast (19).

### Normalization

Features were merged to the genus level by averaging over all features assigned to the same genus. Given the large variation in feature values, we transformed these values to Z scores by adding a small value to each feature level (0.1) and calculating the 10-basis log of each value. Statistical Whitening was then performed on the table, by removing the average and dividing by the standard deviation of each feature.

### Machine Learning

Supervised Learning was performed on the normalized and merged version of the 16S rRNA feature table to recognize patterns in the data. Principal Component Analysis (PCA) was performed using Python version 3.5 and its package sklearn. A 2-tailed p-value of less than 0.05 was considered to indicate statistical significance. A LASSO regression was performed over the projection of the normalized features on the Primary Components (PC) of the PCA to predict future BMI change. Leave One Out cross-validation method was performed. More complex methods were not used to limit overfitting, given the limited number of samples.

### Statistical analysis

All correlations studied here are Spearman correlations. P values of ROC curves are computed using scrambling the classes (positives or negative) of the samples and computing the AUC of 1,000 scrambles. The real AUC was compared to the 1,000 scrambles. Benjamini Hochberg correction was performed when multiple correlations were computed for each PC (for example when correlating age with PCs of microbiome projection).

## Results and discussion

To test the possibility of predicting bariatric surgery outcome, we analyzed 265 fecal samples from patients of 2 main groups: obese who underwent bariatric surgery and naturally slim. For the obese patients (BMI>35) we sampled the microbiome at five time-points (Supp. Mat Fig 1) – one at enrollment (A, 66 samples), three weeks after a low carbohydrate diet and immediately before the operation (B, 58 samples), and three time-points following the surgery (two weeks– C, 23 samples; three months – D, 22 samples; and six months E, 9 samples). Not all individuals have been sampled at all time points. This was compared to 83 slim control individuals (BMI 19-25) (For all details, see Methods). We collected BMI and sugar A1C information for the same patients in late time points up to a year and a half post-surgery to track their weight loss and the remission of diabetes. The obese passed Sleeve, Omega Loop and Roux-En-Y surgeries, with an approximately equal fraction (Fig 1).

**Figure 1.**
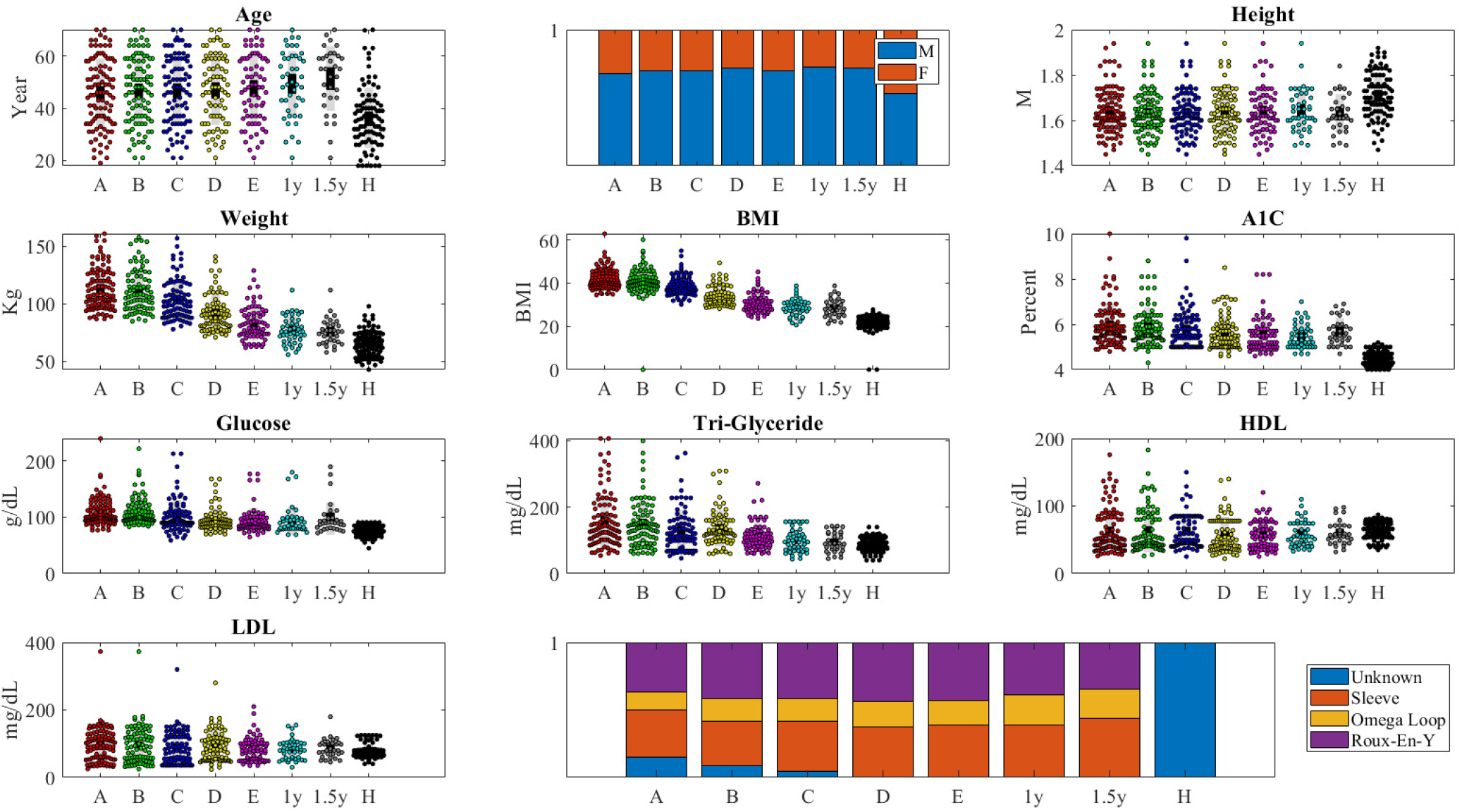
Distribution of Age, Gender, Height, Weight, BMI, A1C, Glucose and Tri-glyceride levels, HDL and LDL at the samples taken in each group and time point (H is slim/ All other samples are obese before (A,B) or after surgery (C,D,E,1y,1.5y).

The slim population was younger and had more females compared to the population who underwent surgery (36 +/−12 vs 48+/−12 and 50% males vs 29%). Overall, the patients’ mean BMI was reduced from 43.3+/−6.8 (Mean+/−SD) to 27.8+/−1.5, which represents an average loss of 84.7% overweight (compared to BMI 25), blood sugar levels were reduced from Hemoglobin A1C of 6.5+/−0.4 to 5.8+/−0.75 or from blood sugar levels of 125+/−11 g/dL to 95.4+/−10, Triglyceride levels decreased from 183+/−20 to 102+/−13. All parameters described are significantly (P<0.001) lower from their starting point and not different from the slim control (Fig. 1).

The gut microbiome of all donors was analyzed using 16S rRNA gene sequences (emphasizing the 16SMetaVx.V2) (20), and feature tables were produced using QIIME2 (16). The feature tables were then merged to the genus level. Taxa appearing in less than 5 samples (each sample is one time-point of one host) were removed. All samples were log-normalized to highlight the differences in rare bacteria and z-scored. The normalized tables were projected on PCA components. The first step aim was to homogenize the description level and reduce the dimension. Since multiple features are associated with the same bacteria, and some features are associated with different levels of classification, we averaged all features associated with the same species in each donor (Fig. 2A-C). Note that while information is lost in the process, such a process is essential for the following machine learning. We have previously demonstrated, in multiple microbiome-based machine-learning studies (21–23), that averaging over all features representing the same genus or species improved the prediction accuracy. If a feature was only present in part of the samples, it was given a value of 0 in all other samples. The resulting values have a scale-free distribution, which often masks large changes in relative frequencies of rare bacteria. To handle that, we log-transformed all feature values and added a small constant value (0.1) to avoid log of zero values. This allows for a narrower distribution of values (Fig. 2D). Finally, given the very high correlation between the relative abundance of different bacteria (Fig. 2E), we projected the z-scored bacterial expression level at all time points and in the slim population to principal components, which capture most of the variance in the bacterial diversity (Fig. 2F).

**Figure 2.**
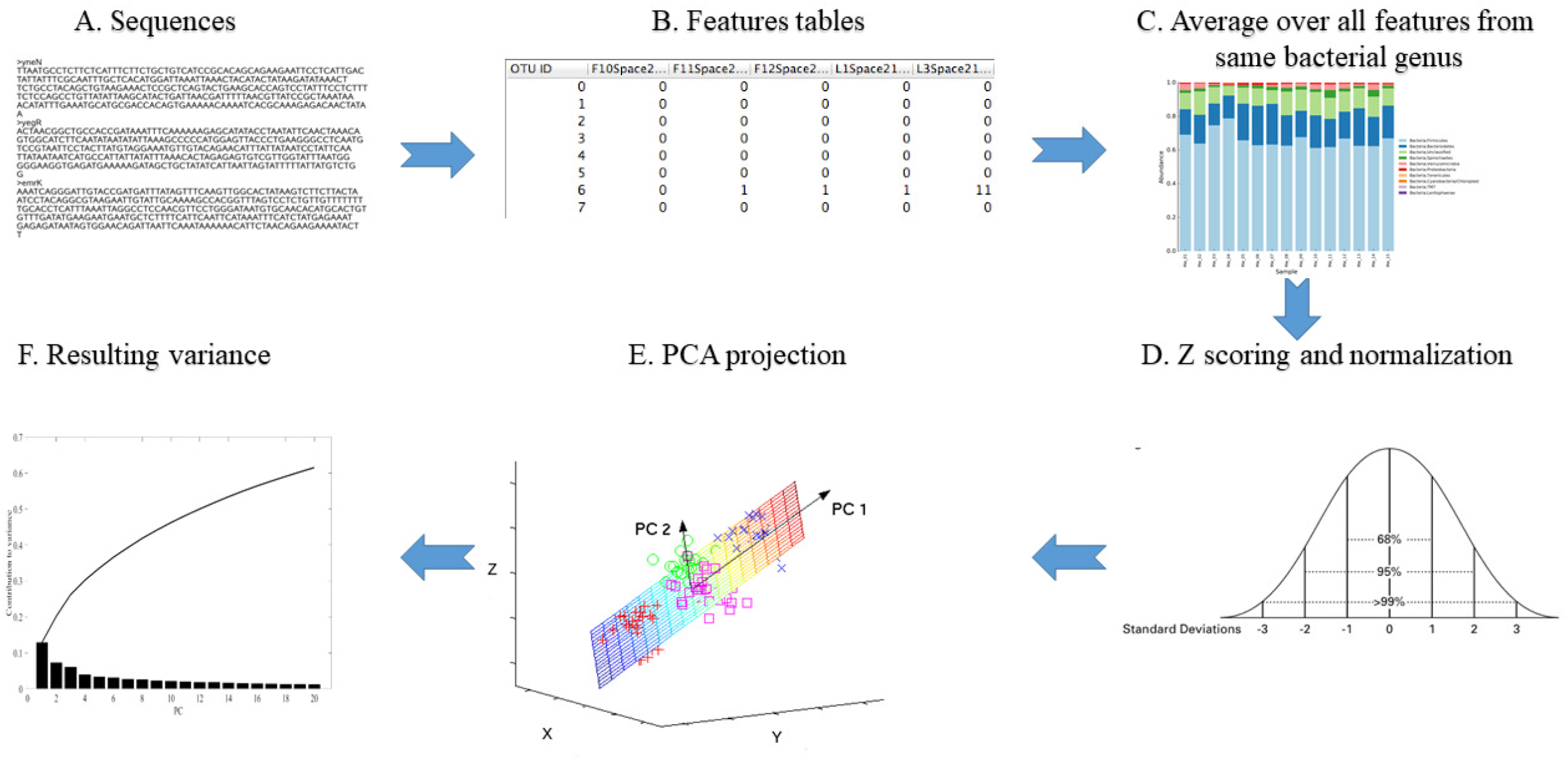
Outline of analysis (from upper left to upper-right and then from lower right to lower left). First, the Fastq sequences are quality controlled. The good quality sequences are translated to features using QIIME2. To homogenize the description level, the feature levels belonging to the same genus in a given sample are averaged to the genus level. (Lower level). The sample distribution is heavy-tailed. It is thus log-transformed with a minimal value (0.1) added to each feature level to avoid log of zero values. The results are then z scored by removing the average and dividing by the standard deviation of each sample. The dimension of the z-scores is further reduced using PCA. The first 8 PCs explain approximately50 % of the total variance (lower left figure).

The projection on the first principal vectors (PC1, PC2, PC4 and PC5) delineate axes separating the obese from the slim individuals (Fig. 3A,B). PC3 showed no correlation with BMI (Fig 3A). The clear separation of the projections on the first PCs agrees with observed major differences in the microbiome of slim and obese individuals. The large BMI difference between groups (BMI>35 in obese versus BMI of <25 in slim) translates to a large difference in the microbiome. We next tested whether diet or bariatric surgery push back the population toward the slim profile. The results are surprisingly opposite (Fig. 3B). The distance between the projection on the first PC of the post-diet and post-surgery and the slim profile keeps increasing and reaches a maximum after a year. The major difference between these projections allows for a simple classification even with a linear SVM of slim vs obese and pre vs post-surgery samples (Fig. 3C,D). The main contributions to both classifiers are from PC1, PC1 to PC5 for the healthy (H) vs obese (O) (Fig. 3E for contribution and Fig. 4A-D for composition). Note that higher test AUC can be obtained by non-linear classifiers. However, the linear classifier gives a clear picture of the contribution of each PC to the microbiome development.

**Figure 3.**
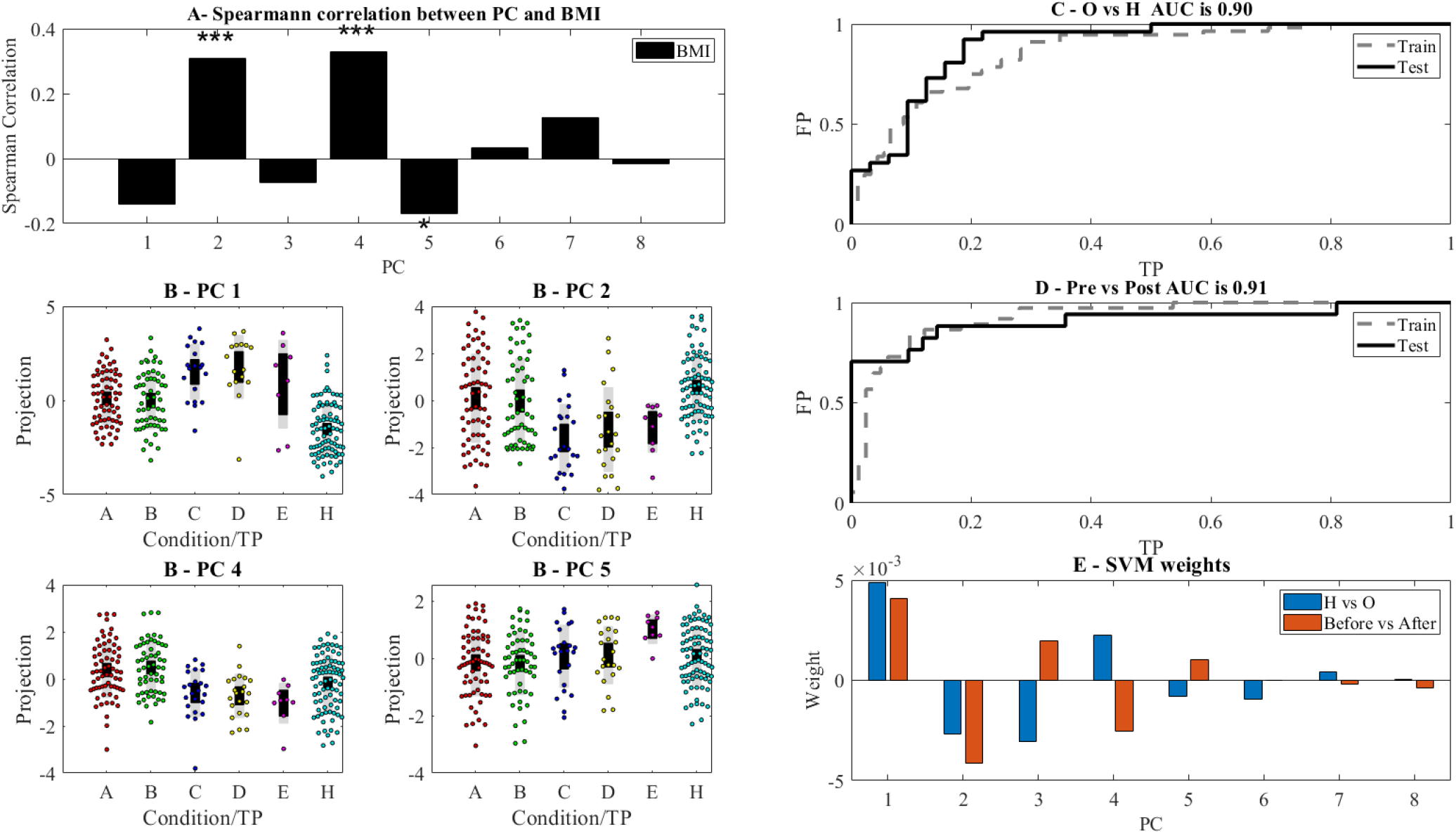
A - Spearman correlation between BMI of samples and the eight highest variance PC. ***,** and * represent significance level of 0.001,0.01 and 0,05 respectively (in this and all following figures). PC2,4 and 5 are the most correlated with BMI.B - Projection of the significant PC on the different stages. One can see in all PC a clear difference between the H and the obese states. Following surgery, the projection is farther away from the H state than before. Note that we do not have microbiome samples from the latest time points. C,D ROC curves of linear SVM classification using the projections on the first 8 PC of the H vs O and within the O group before vs after surgery, and (E) the resulting weights (lower right plot)

**Figure 4.**
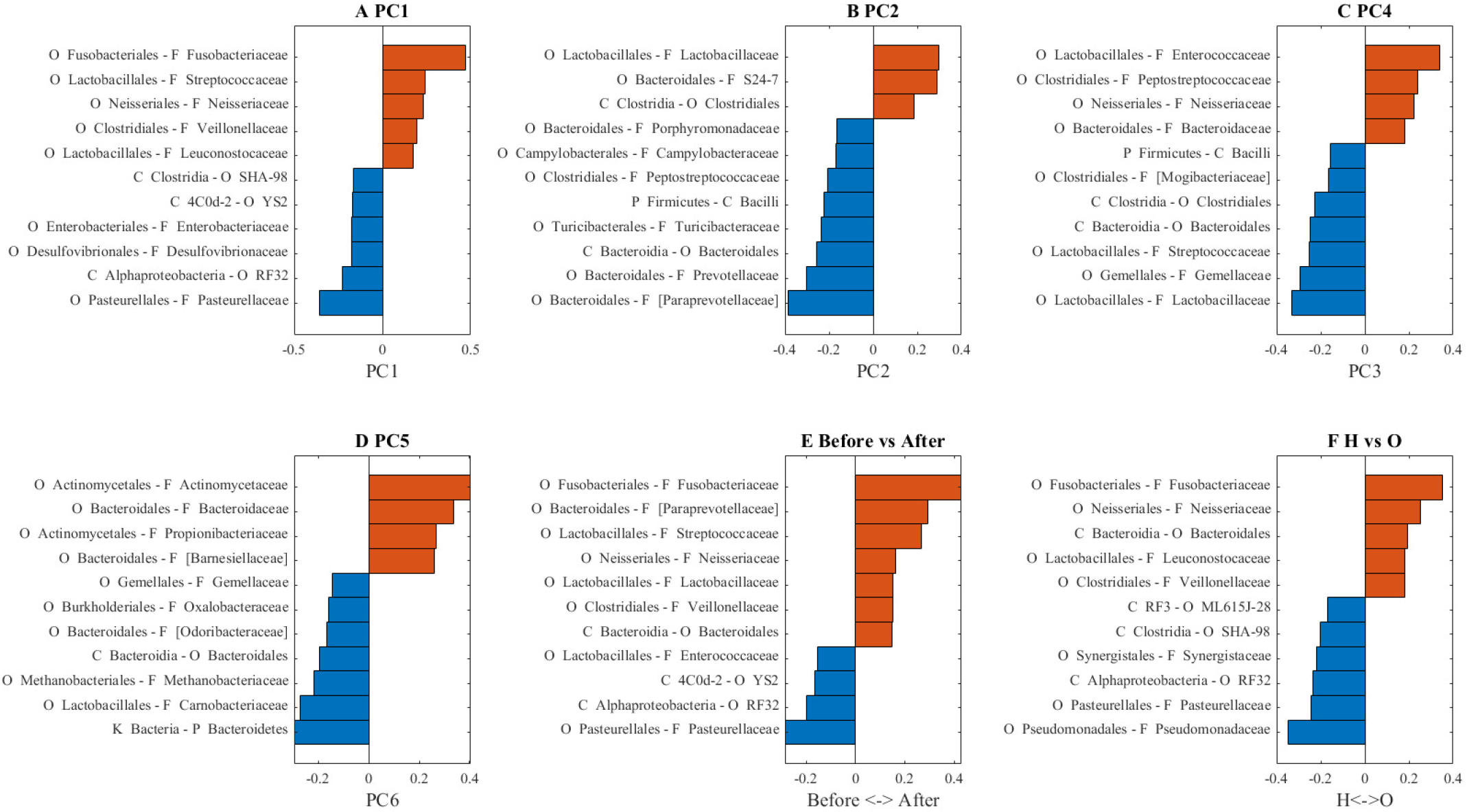
A-D Bacterial composition of PC1,2,4 and 5 E,F Weights of linear SVM classifier for H vs O and before vs after surgery. Only the top 15 % of features are presented (based on their absolute weights).

One can then project back the correlations between the PC and the state/BMI to the original features, and find features that are correlated to BMI (PC 1,2,4 and 5; Fig. 4A-D, respectively), the features that change significantly after surgery compared to before surgery (Fig. 4E) and the features that are over and underrepresented in obese individuals compared to healthy (Fig 4F).

To test for possible confounding effects, we tested whether the observed changes in the profile may be the result of age or gender, or whether they are related to the total BMI. There is a limited correlation with age and no significant correlations with gender and age of all other PC (Fig. 5A,B).

**Figure 5.**
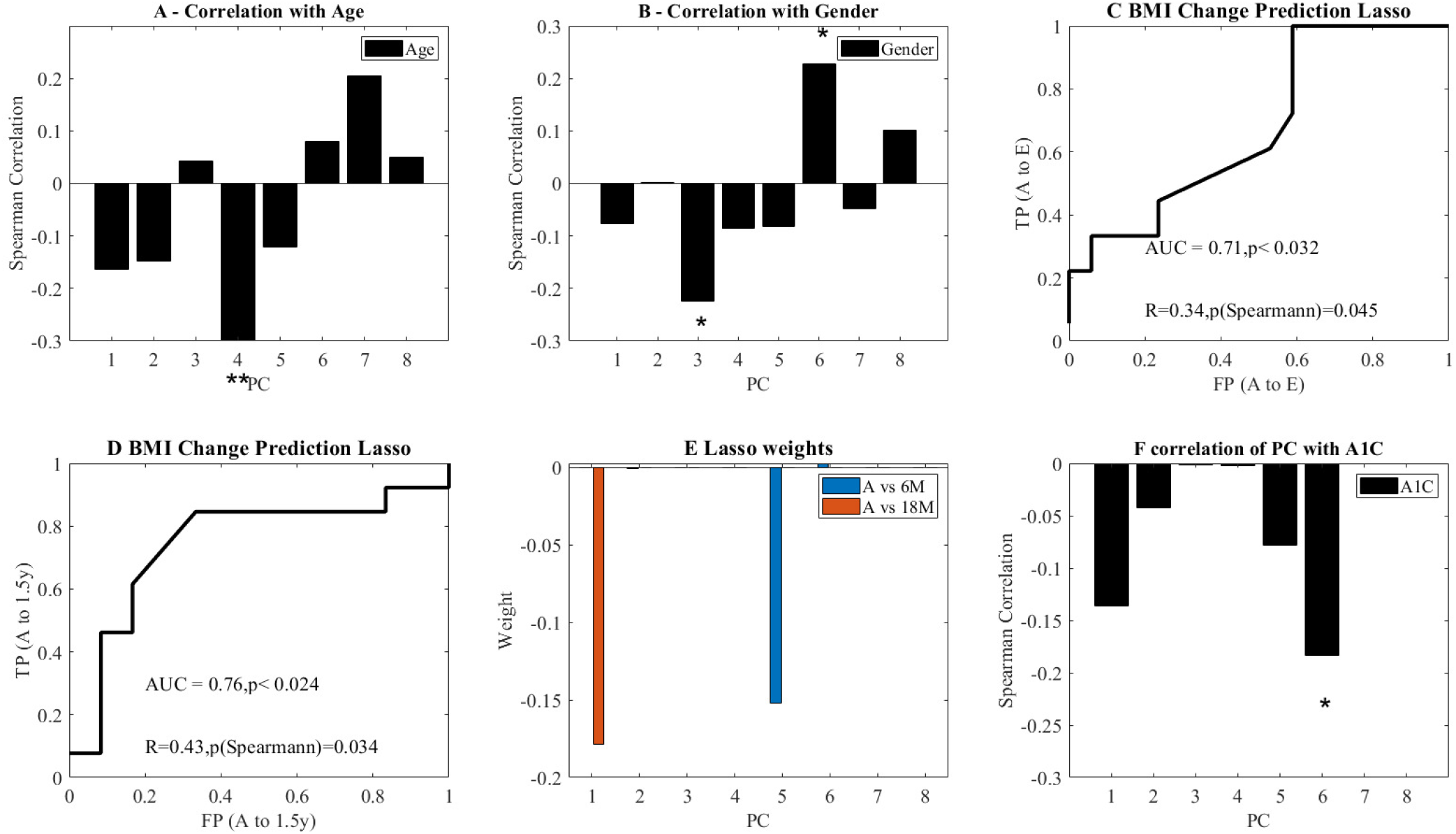
A,B correlation of Age, Gender (represented as a one-hot) and BMI with the PC. C-D ROC curves of future BMI change based only on the PC at point A. The binary predicted change is above or bellow median change. The ROC curves for above and below median change in BMI, using a LASSO regression and a LOO validation. C is for six months and D is for 18 months. The first p-value is the ROC curve p-value compared with AUC of 1000 scrambling of the predicted values. The second p is the Spearman correlation p-value between predicted and actual change. E - Average regression weights over all LOO learning sessions. The only non-zero coefficients are the 5thPC for 18M and the 1st PC for the 6M B-F Average weights of coefficients in the LASSO regression for plots C and D over all LOO predictions.

We then tested whether the same decomposition can be used to predict a future change in BMI. We performed an L1 (Lasso) regression of the projection on the first PCs of the A point and the change in BMI between point A and 6/18 months after surgery. The prediction was tested using a Leave One Out (LOO) methodology, and the Spearman correlations on the test values between the predicted change and the observed change (on the LOO test)), as well as the Area Under Curve of a predictor of whether a patient will have a more/less than average reduction in BMI. The AUC and correlations are significant for both points (Fig. 5C,D). The AUC and correlation for 12 M are also above random, but do not reach significance (data not shown). When looking at which features are contributing to weight loss at 6 months, PC1 and PC5 were the only contributors (Fig. 5E). Note that the prediction is based on the microbiome pre-surgery, so that these results cannot be an effect of surgery type.

Another possible candidate to affect the microbiome is the sugar level. We tested the correlation between the projections on the PCA and the A1C. Indeed, a negative correlation is found between A1C and PC6 (Fig. 5F). This correlation might be used for predicting the disease in healthy subjects both from risk groups and in general.

## Conclusions

To summarize, we have shown that the decomposition of the relative bacteria frequency (as represented by log value) represents different aspects of the donors. This decomposition highlights that people who lost weight after bariatric surgery have a very different microbiome composition compared to people who are “naturally” slim. Furthermore, the more weight they lose, the more their microbiome profile differs not only from their starting profile as obese but also from naturally slim people.

Moreover, one can predict in advance whether surgery will succeed in reducing BMI and whether a subject has diabetes. The PCs are determined by the composition of the studied populations, and the analysis of different populations may highlight different possible projections of the microbiome composition. Interestingly, 2 main PCs remain with no clear correlation to the phenotypes studied here. Those may represent other important elements affecting and affected by the microbiome not tested in this study.

As expected, when comparing the microbiome before and after surgery (Fig. 4E) we found that the three top features which were overrepresented in individuals before surgery belonged to the family Pasteurellaceae and the order RF32 (both part of the Proteobacteria phylum). Proteobacteria have long been correlated with dysbiosis leading to inflammation and obesity (24–26). Some of the mechanisms proposed for the contribution of members of the Proteobacteria to obesity in mice and humans include induction of inflammation and intestinal barrier dysfunction (24). We next compared the microbiota of obese vs. Healthy individuals (Fig. 4F). Here we did not expect to see the same results seen when comparing before vs. after surgery microbiomes as we demonstrated that the microbiome after surgery is not similar to the healthy microbiome. Indeed, the obese individuals had an over-representation of members of the Fusobacteriaceae. Fusobacteriaceae have been reported to directly correlated to gut health and have been shown to increase in multiple inflammatory diseases (27). One of the members of Fusobacteria is *Fusobacterium* which is known for it’s immune modulatory characteristics which eventually render antitumor immune cells inactive (28).

## Declarations

## Acknowledgments

Diana Bluvshtein, Manar Hadad, Natali Ogo for their help as clinical nurse involved in data collection and patient enrolment.

## Ethics approval and consent to participate

Kaplan Medical center 0068-15-KMC

Rabin Medical Center 0088-16-RMC

Tel Aviv Medical Center 0548-16-TLV

Poria Medical center 0057-18-POR

## Consent for publication

All co-authors have agreed to the publication

## Availability of data and material

The Sequence files are now uploaded to the EBI

## Funding

US-Israel bi-national Research and Development Fund (FIRD-F) project # 1459 (RC, LMS, HM, CL)

16S data has been deposited at EBI with Accession number ERP122895

OTU tables are available as Supp. Mat.

## Supplemental material legend

Supp. Mat. Figure 1 – Experimental setup. The experiment tested the microbiome and different intakes (HDL, LDL, Triglycerides, BMI and A1C) of patients from two groups – obese who underwent Bariatric surgery and slim individuals. The slim individuals have been sampled once while the obese patients were sampled in 5 time points –before the entire process (A), after low carbohydrate diet and before surgery (B), 2-3 weeks after surgery (C) 3 months after surgery (D) and 6 months after surgery (E). We further sampled the weight after 12 and 18 months.

Supp Mat.2 All OTUs and taxonomy for all samples used here.

